# Age-Dependent Patterns of Gut *Bifidobacterium* Microbiome Diversity across child Cohorts in a Multi-Ethnic population, Western China

**DOI:** 10.1101/2025.02.10.637477

**Authors:** Jian Huang, Yuxin Chu, Liang Zhixuan, Huimin Zhang, Quanhao Zhao, Fengwei Tian, Yongqing Ni

## Abstract

High-throughput sequencing has provided unprecedented opportunities to elucidate the role of human gut microbiota. However, current studies have primarily focused on infant and adult cohorts, leaving biomarkers associated with the development of transitional children unclear, particularly in revealing geographic, multi-ethnic group underrepresented. Here, we analyzed fecal samples from 776 infants and children aged 0 to 15 years with amplicon sequencing of the 16S rRNA and *groEL* gene, representing multi-ethnic groups from different residential locations in the Yining region of Xinjiang. First, we systematically characterized the gut microbiota Spectrum of children in the Ili region. As age increases, the diversity and functional composition of the microbiota become richer and more stable, with *Bifidobacterium* consistently dominating the community. Utilizing Bayesian regression models, we identified a cluster of microbiota strongly associated with growth and development in children, which phylogenetically corresponds to *Bifidobacterium breve*. Finally, we included another similar study in the species-level of *Bifidobacterium*, and metrics of the age-related effect sizes showed heterogeneity across ethnic groups (Uyghur vs Han: β_Deming_ = 4.40 and Kazakh vs Han: β_Deming_ = 4.33). In conclusion, this study provides novel insights into the developmental trajectories of the gut microbiome in children and its relationship with health.

**IMPORTANCE:** The intestinal microbial markers associated with the development of geographically diverse, multi-ethnic children, particularly those represented by Bifidobacterium species, have been elucidated in this study. Through the application of Bayesian regression models, we identified a microbiome profile closely linked to childhood growth and development, specifically phylogenetically aligned with *Bifidobacterium breve*. When incorporating a comparable study conducted at the species level of another *Bifidobacterium*, it was observed that age-related effect sizes varied among different ethnic groups, offering novel insights into the developmental trajectory of children’s gut microbiota and its implications for health.

## Introduction

The human gut contains trillions of microorganisms whose composition is related to dietary preferences, individual genetics, and innate immunity, and together they form a complex ecosystem of the gut. (1-3) The gut microbiota plays a crucial role in the human environment. It evolved alongside its host to perform essential physiological functions for the host, such as preventing infection by various pathogens.(4, 5) Despite the acknowledged relationship between microbiome composition and health, the scientific community persists in the quest for microbiome biomarkers suitable for disease diagnosis and clinical applications.(5, 6) However, variations in microbial composition among individuals(7, 8) and the impact of environmental factors(9) on microflora shifts within the generally population significantly hinder this research. Current research is predominantly centered on highly developed countries,(10) resulting in a lack of diversity in study populations that constrains the generalizability of findings to broader global contexts.

Moreover, the development and maturation of the gut microbiota represent a highly dynamic process that begins at the newborn stage, influenced by factors such as maternal microbiota(11) and the mode of delivery.(12, 13) In fact, a considerable part of members was introduced into the human body through environmental exposure and diet, not persistent gut residents. (14-16) Instead, only a subset of gut microbes, primarily composed of obligate anaerobes, establish mutually consistent and stable symbiotic relationships with their hosts.(17, 18) As a result, the spatiotemporal dynamics of the microbiome may not invariably lead to the formation of stable microbial communities within the first 3 years of life,(14, 19) but instead continue to evolve until middle childhood or even later.(19, 20)

Bifidobacteria are a paragon of symbiotic bacteria in the human gut, particularly during early life. (21-23) *B*. *longum* is globally the most prevalent subspecies irrespective of age, and its strain transmission is commonly observed within family or even between unrelated subjects. (24, 25) Prevalence of other species in adults, such as *B. bifidum* (26) and *B. breve* (27), varies widely across cohorts and individuals.(18) Numerous studies have demonstrated a positive correlation between *Bifidobacterium* and various physiological functions of the human host throughout the lifespan.(28, 29) In general, Bifidobacteria, coevolved with their hosts, could be categorised into two main groups based on their origins: Human-Residential Bifidobacteria (HRB), which are naturally found in the human gastrointestinal tract and believed to have coevolved with their hosts, and non-HRB species that inhabit animals or the environment.(28, 30) Nowadays, what we do know is that some HRB are highly abundant colonizers in the early stages of human life, and their presence is associated with many beneficial host effects, even last a lifetime. The abnormity of HRB may increase the risk of disease in childhood or later in life.(31)

In fact, *Bifidobacterium* is not the dominant bacteria in the adult gut compared to the infant gut microbiota, with a relative abundance of only 1% or less, and even undetected in the gut of some populations.(32) Particularly, the pitfalls inherent to 16S gene region or subregions (the V4 region or the V3-V4 region), including the limited discriminating power among sequences and the under-representation of reads belonging to bifidobacterial taxa, may conceal the discovery of species- and strain-level sequence variants,(33, 34) leading to skewed estimates of bifidobacterial community. Overall, although early life-associated dynamics and diversity of *Bifidobacterium* species have been well studied, there is limited understanding of the diversity of *Bifidobacterium* at the species level across different human populations.(35) During the early stages of human development, the ecological significance of species and strain diversity within the gut *Bifidobacterium* microbiome, as well as age-dependent growth patterns across different human populations, warrant further investigation.(7) Access to extensive cross-sectional data sets across diverse populations is expected to facilitate a more in-depth exploration of specific strains and species of *Bifidobacterium* in the gut microbiome, as a function of child age, lifestyle, and health status.

The gut microbiome composition significantly impacts health, especially during childhood, a vulnerable period linked to ecological imbalances and diseases. Therefore, from the perspective of diverse populations, elucidating the composition of intestinal microflora in early childhood is crucial for the development of new therapeutics and for fostering the growth and development of children. In this study, we used metataxonomic data from the 16S rRNA gene for bacteria, along with *groEL* gene sequences specific to *Bifidobacterium*, to analyze the gut microbiomes of 776 children aged from newborn to 15 years old in Yili prefecture, Xinjiang, western China. This area is known for its diverse ethnic groups and limited inter-ethnic marriages. This sympatric human populations with different lifestyles are good models to investigate how genetically relatively isolation can shape microbiome diversity patterns. Our aim was to gain insight into the diversity and composition of bifidobacteria community in cross-sectional children‘s cohorts across different ethnic groups, and to in-depth explore specific strains (species) of *Bifidobacterium* in the gut microbiome as a microbial signature indicative of child age, ethnicity, and body mass index.

## Result

### Description of study population

We ultimately had a total of 776 children aged from newborn to 15 years old (Table S1 and S2), who reside in a relatively concentrated manner, exhibit limited mobility, and demonstrate pronounced ethnic representation. Participants recruited participants from Ili Kazak Autonomous Prefecture in Xinjiang, China. The participants were unevenly distributed across ethnicity, due to the ethnically diverse and locally concentrated nature of the Ili region. *Kashgar County* is predominantly inhabited by Kazakhs, while *Yining County* is primarily populated by Uyghurs. In contrast, other regions (*Nilka County* and *Sunzhaqi Niulu Village*) included in our study demonstrate a more ethnically pluralistic composition. The children were initially split into three age groups: Infant group (about 0 to 3 years), kindergarten pupil group (about 4 to 7 years) and primary school pupil group (about 7 to 15 years) at the time of enrolment. The mean age of the participants was 7.32 ± 4.05, and the distribution was roughly uniform. Notably, the BMI z-scores of our randomized recruits exceeded the baseline set by the World Health Organization (WHO) (Supplementary Figure S1).

### Gut microbial characterization

We conducted amplicon-based sequencing of the V1V3 region of the 16S rRNA gene on 776 fecal samples. Sequence analysis using the EasyAmplicon pipeline(36) produced 53,544,690 high-quality paired readings, and the average sequencing depth of each sample was 34,411 sequences (range was 12,845-201,871). We obtained 34,395 high-fidelity zero-radius operational taxonomic units (ZOTUs) in total. Rarefaction curves indicated Observed ZOTUs reached saturation in all samples (Fig 1b). Characterization of the phylogenetic variation showed 9 phyla, 18 classes, 32 orders and 67 families, spanning across 119 genera (Fig. 1c). Most of them belonged to Firmicutes (64.4%), followed by Actinobacteria (17.4%), Bacteroidetes (8.8%) and Proteobacteria (6.9%). To analyze the gut microbial composition of different sampling areas and age groups, we divided them into nine age-level groups (as detailed in the second layer of Fig. 1a), and leveraged an UpSet plot (Fig. 1d) to count the shared microbial ZOTUs. Among the nine groups in our study, the city infant group (blue dot) exhibited a notably low diversity of ZOTUs, while the other eight groups displayed the highest number of unique ZOTUs. The infant groups (red line) shared a high number of microbial ZOTUs. Among these, the Nilka County infant group, with 1,143 ZOTUs, exhibited a notably high number of unique ZOTUs. In addition, we found that the number of common microbes shared among all groups ranked third in the UpSet plot, demonstrating that despite some distinctions, the gut microbiota composition of children in the Ili region remains largely similar. Moreover, we define the core microbiota as the bacterial taxa present in over 90% of the individuals studied. This core microbiota includes nine genera belonging to two phyla: Actinomycetota and Bacillota (Fig. 1e). The core microbiota includes nine genera belonging to two phyla: Actinomycetota and Bacillota (Fig. 1e). Only four genera were previously identified as a global cohort(37), highlighting the distinct nature of the gut microbiome composition in regional populations of children compared to global populations.

**Figure 1.**
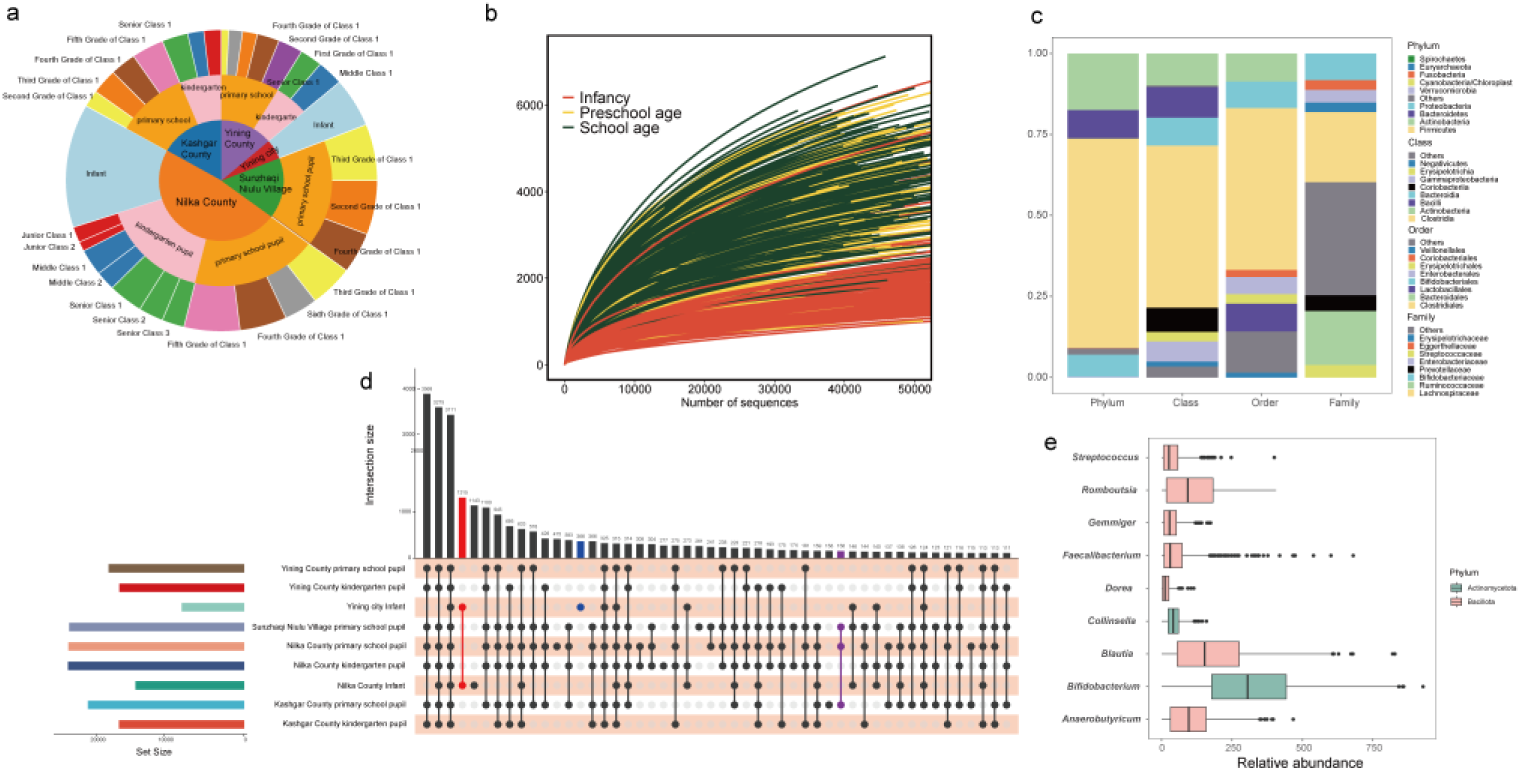
Gut microbial characterization in Ili region children. (a) In this study, the sampling sites of the three-level classification label pie chart. (b) Rarefaction curves of ZOTU measures, with each line representing a single sample. (c) Taxonomic distribution of the Ili region children fecal microflora at phylum, class, order and family levels. Only the top eight taxa are shown at each taxonomic level, and the remaining ones are grouped as ‘others’. (d) The UpSet plot is a visualisation of the overlapping ZOTU clusters across the nine sampling positions. The chart at the bottom represents the various intersections among different categories, with each category depicted as a dot and intersecting categories connected by straight lines. The sizes of the intersection sets are shown in the vertical bar chart. (e) Unweighted mean relative abundance of core genera across the entire dataset.

### Microbial communities in both diversity and composition in different developmental stages

To elucidate the distinct microbial communities associated with different developmental stages of children in terms of both diversity and composition, we compared the microbial relative abundance (Fig. 2). Infants consistently exhibited lower alpha diversity (Wilcoxon test, P < 0.001) compared to older age groups (Fig. 2a, Fig. S2). In contrast, no significant differences in diversity were observed between the school-age and preschool-age groups. Subsequently, we categorized the samples based on grade level and various sampling sites; however, no significant differences were observed as the samples progressed toward a (pre)school-age gut microbiota (Fig. 2b). This may be attributed to the fact that, after the age of 3, children’s gut microbiota typically reach a state of relative maturity, exhibiting minimal changes thereafter(38). Principal coordinate analysis (PCoA) based on Bray-Curtis dissimilarity revealed distinct clustering of samples from children at three developmental stages. Significant differences were observed between infants and both preschool-age and school-age stages (adonis PERMANOVA test, *P* < 0.001, permutations = 1000). Additionally, a significant difference was noted between early childhood and primary school (F = 7.87, *P* < 0.001). The composition of intestinal flora was clearly associated with the developmental stage of the children (Figure 2c). Among the genera with a prevalence ranked within the top 11 and a mean relative abundance exceeding 1% (Fig. 2d), *Bifidobacterium* (61.58%), *Streptococcus* (9.13%), and *Collinsella* (4.53%) were predominantly characterized by their high abundance during infancy. As children transition into toddlerhood, the dominance of *Bifidobacterium* declines to 28.31%, with *Anaerobutyricum* (6.48%) and *Romboutsia* (5.48%) emerging, indicating a significant increase in microbial diversity. By elementary school age, *Bifidobacterium* levels rebound to 32.02%, while *Romboutsia* (6.58%) and *Blautia* (5.57%) remain prevalent. At the same age and in different populations, the composition of microbial genera is generally similar (Fig. 2e).

**Figure 2.**
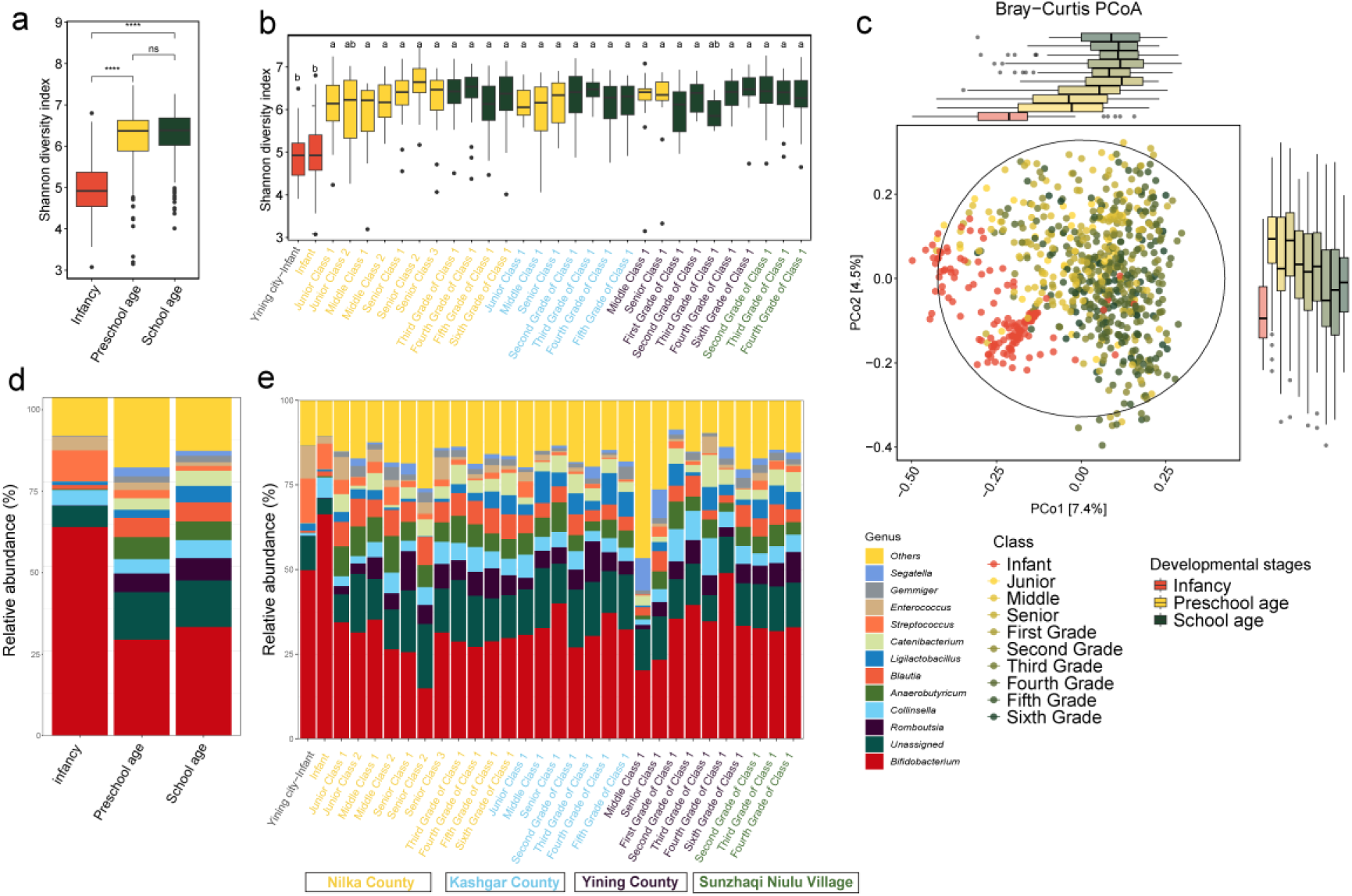
Distinct diversity and composition of gut microbiota from different developmental stages of children. (a) Alpha diversity (Shannon diversity index) of infants (n = 145), preschool children (n = 212) and school-age children (n = 421). The p-value was computed using “ggpubr” packages based on the method = “wilcox.test” (**** *P*<0.0001, ns: no significance). (b) Alpha diversity (Shannon diversity index) of stools from developmental stages children stratified by “sampling class and position points” into five categories for position points and thirty categories for class points. The p-values were computed using the Wilcoxon test. The p-value was calculated with the “dunnTest” function from the FSA package, applying the Benjamini-Hochberg method for multiple comparison adjustments, and are represented by lowercase letters (*P* < 0.05). (c) Principal coordinate analysis (PCoA) of samples of different developmental stages of children was conducted based on Bray-Curtis dissimilarity of species. The boxplots on the right side and below show samples of different age-stages projected onto the first two principal coordinates, respectively. The p-values were calculated by the adonis function (permutations = 1000) function from the R package “vegan” for the PCoA plot. (d, e) Comparisons of the mean relative abundance of gut genera from three developmental stages of children (d), and stratified by thirty class points (e). Only the top 10 abundance genera were displayed. The X-axis is distinguished by colour-coded labels, denoting distinct sampling locations.

### Differences of gut microbiota in three developmental stages of children

To better understand enhance comprehension of the developmental trajectory in fluences bacterial community composition, ZOTUs that are specifically enriched in the three developmental stages of children were identified. Using the LEfSe analysis, the values of Linear Discriminant Analysis (LDA) for various species were calculated across different developmental periods, and species with LDA values exceeding 2 were identified as significantly different. The gut microbiota is characterized by a significant number of ZOTUs enriched in different age groups: 1214 infant-enriched ZOTUs, primarily from *Bifidobacterium*, *Catenibacterium*, and *Anaerobutyricum*; 312 school-age-enriched ZOTUs, mainly from *Romboutsia*, *Bifidobacterium*, and *Blautia*; and 113 school-age-enriched ZOTUs, predominantly from *Bifidobacterium*, *Intestinibacter*, and *Anaerobutyricum* (Fig. 3a and Supplementary Table S3).

**Figure 3.**
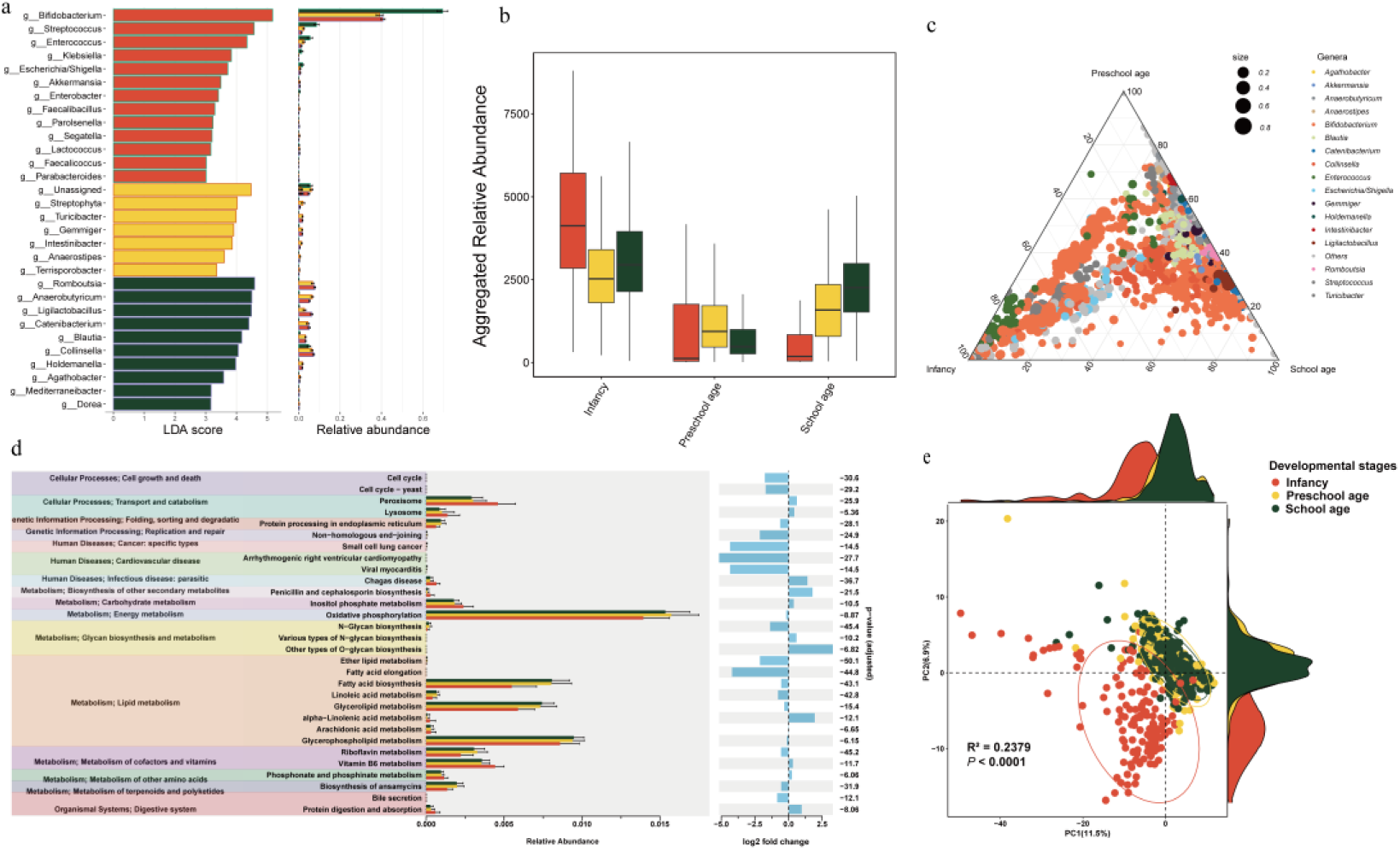
Differences of gut microbiota across three developmental stages in children. (a) LEfSe analysis of microbial profiles between infancy and (pre) school-age at the genus levels. (b) Aggregated relative abundances of each group of enriched ZOTUs (infancy-, preschool age- and school age-enriched ZOTUs) in each compartment for the stool sample. In each compartment, the difference from 100% RA is explained by ZOTUs that are not significantly enriched in a specific compartment. (c) Ternary plots depicting compartment relative abundances of all ZOTUs (> 0.1 ‰) for the three developmental stages of children. (d) Predicted functional analysis of stool samples revealed pathways affected by the age development. Predicted functional pathways were identified using PICRUSt, and the differential abundance of these pathways is presented. (e) Principal components analysis (PCA) was conducted based on predicted functional pathways of intestinal microbiota from the three developmental stages of children, assessed by PERMANOVA. Ellipses represent 95% CI.

The characteristics of the bacterial community are further elucidated (Figure 3b), which presents the aggregated relative abundances (RAs) of ZOTUs specifically enriched at a particular developmental stage. As anticipated, this analysis revealed a decreasing contribution of the infant-enriched ZOTUs across samples from infants, preschool, and school-age children, with mean aggregated relative abundances (RAs) of 76.11%, 13.55%, and 10.32%, respectively. No distinct differences in enriched ZOTUs were observed during the preschool stage, and furthermore, the ternary plots also corroborate this phenomenon (Fig. 3c). This observation appears reasonable, as childhood development represents a transitional stage that may not result in a enriched gut microbiota.

To assess potential differences in the functional composition aspect of the intestinal microbiome, we conducted PICRUSt2 analyses across the three developmental stages. The results revealed a significant difference across the predicted KEGG pathways at (pre)school age compared with the infant stage (Fig. 3d). PICRUSt2 analysis revealed that the primary functions of intestinal microbes in children from Yining, were predominantly associated with protein families, genetic information processing, signal transduction, cellular processes, and carbohydrate metabolism. Additionally, pathways related to human diseases and polysaccharide synthesis were significantly elevated compared to the infant group (*P* < 5 × 10^-8^, and |log2 fold change| > 2). Fig. 4d illustrated the principal component analysis (PCA) of the predicted intestinal microbial functions across the three developmental stages. We observed significant differences in intestinal function between the infant group and the other two groups (*P* < 0.0001). The infant group exhibited considerable heterogeneity, while the preschool age group demonstrated homogeneity in predicted intestinal microbial functions. This may indicate that the functions of intestinal microorganisms tend to become more concentrated and stable as children progress through their developmental stages.

**Figure 4.**
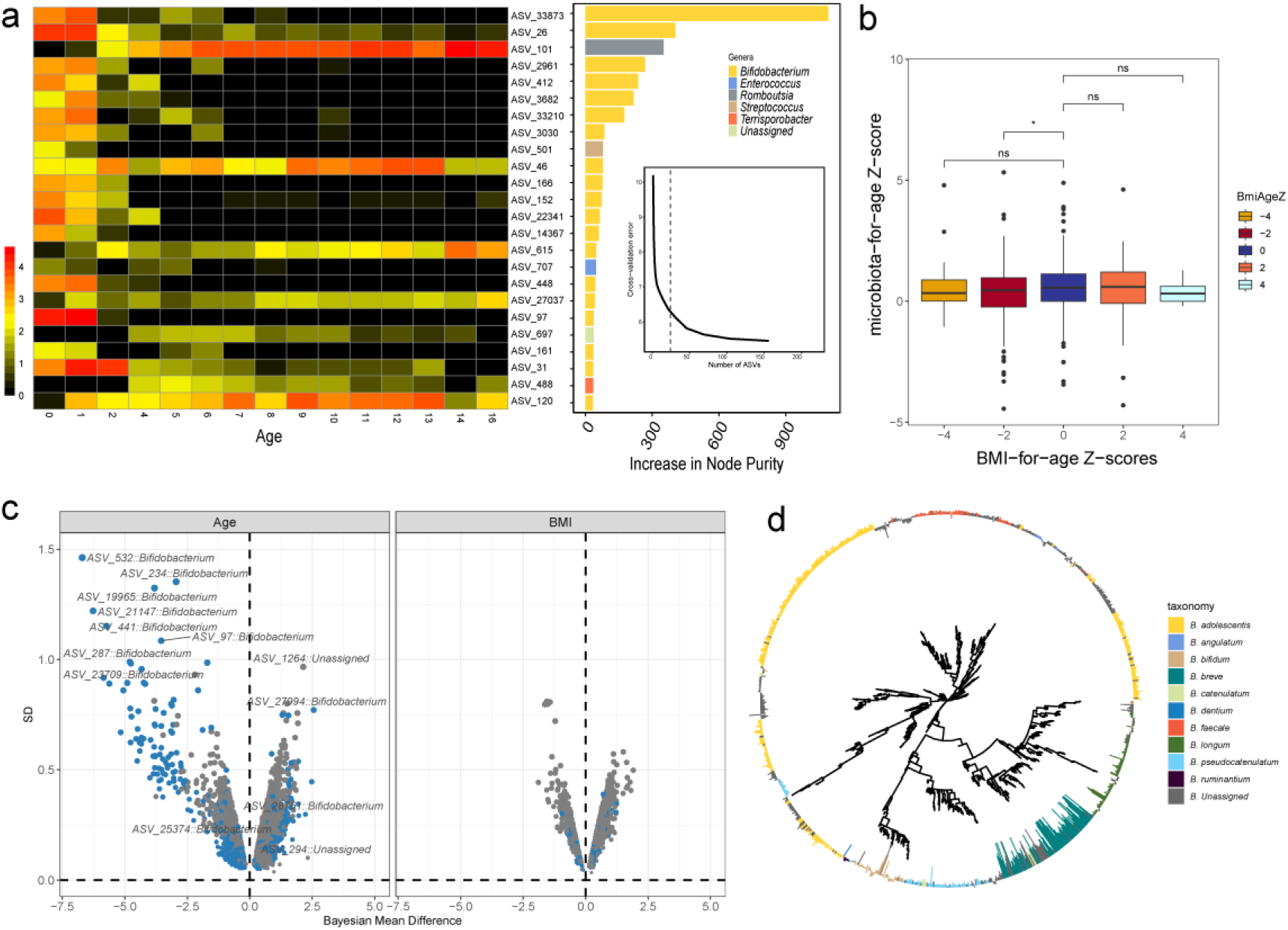
Gut microbiota maturation obtained from concordant healthy children. (a) The Random Forest regression analysis of fecal bacterial ZOTUs from a training cohort of healthy Yili infants and children (n = 526, BMI Z-scores: [-2, 2]) in relation to age-discriminatory has identified a ranked order of taxa with significant age-discriminatory potential. A heatmap illustrates the temporal dynamics in the relative abundances of the 24 most discriminatory ZOTUs within the fecal microbiota. These taxa, ranked by their feature importance (mean ± SD of the Increase in Node Purity), are accompanied by their respective taxonomic classifications. (b) Microbiota-for-age Z-score in children with different health conditions. Using samples with a BMI ranging from -2 to 2 as the training set, the relative microbial maturity of children was predicted based on their BMI categorisation into the following groups: -2 to -4, less than -4, 2 to 4, and greater than 4. (c) Bayesian effect size analysis was conducted to evaluate the impact of age and BMI on the abundance of taxa. The graph plots the influence of age and BMI on taxa, with the effect size represented on the X-axis and the error margin displayed on the Y-axis; Attributes of taxonomic data (ZOUTs), only when show if the 95% confidence interval for any of their age or BMI effect sizes did not contact the 0 coordinate. (d) Age-related bifidobacterium phylogenetic tree. The GTR model was constructed based on 16S rRNA to draw the phylogenetic tree. With the color of the outer ring representing species-level bifidobacterium species annotation, and the concave bar chart having a negative effect and the convex one having a positive effect.

### Gut microbiota development trajectory in growing children

To explore the potential causal relationship between microbiota maturity and child development trajectories, we identified age-discriminatory taxa through Random Forest regression analysis. The maturation of microbiota was modelled in 526 healthy infants and children (BMI Z-scores: [-2, 2]) through the regression of ZOTUs against chronological age, utilising this as the training set (Fig. 4a). To ascertain the validity of the model, we employed the 10-fold cross-validation method to evaluate its predictive performance, revealing 69.14% of the variation associated with the age factor, which demonstrated a positive and significant correlation between the host’s chronological age and the predicted microbiota age (Fig. S3, Fig. 4a).

As Subramanian(39) proposed, the relative maturity of the microbiota was assessed by Z-scoring the predicted ages, resulting in the microbiota-for-age Z-score (MAZ) for each group. MAZ provides insights into the variations in microbiota age predictions based on differing host age ranges. However, our analysis revealed no significant differences in MAZ between the normal group and either the obese or lean groups (Fig. 4b), a finding that contradicts the conclusions of previous studies. This discrepancy may arise from the fact that the study participants were individuals who self-reported as healthy and did not exhibit any developmental diseases.

In light of the aforementioned findings, we carefully investigate the properties of the microbial count data, a thorough investigation was conducted into the properties of the microbial count data, and we calculated the dispersion degree and zero expansion probability of the data from different classification levels. We constructed a zero-inflation Bayesian regression model(40) to measure the effect of each taxon on age and BMI. Figure 4c shows the size of the effect size of ZOTU covariates with age and BMI differentiation. A total of 5243 ZOTUs were identified where the effect size and 95% confidence interval did not overlap with the zero scale, and 1847 for BMI. Among these, 1,053 exhibited positive effects of the age factor while 153 negative effect sizes with absolute values greater than 2.5, along with 5 additional positive effects exceeding this threshold. We did not find ZOTUs with absolute effect sizes greater than 2.5 for BMI. The genera associated with positive age benefits include *Parabacteroides*, *Blautia*, and *Faecalibacterium*. Notably, among the ZOTUs with absolute negative age effects greater than 5, bifidobacteria were predominantly represented. This implies a strong reduction in bifidobacteria during the child’s growth and development trajectory.

To further investigate the age-related phylogeny of *Bifidobacterium*, we constructed phylogenetic trees for Bifidobacterium with significant effects, with the outer bar chart illustrating the effect sizes (Fig. 4d). The results demonstrated that ZOTUs with similar evolutionary distance are also exhibit comparable age-related effect sizes. *B. breve*, *B. pseudocatenulatum* and *B. longum* also exhibited a negative correlation with age, while *B. adolescentis* and *B. faecale* showed a positive correlation with age. *B. bifidum* demonstrated both positive and negative age-related ZOTUs, however the smaller classification (smaller than the species level) showed consistency and did not rule out the cause of amplification bias.

### Gut Bifidobacterium spectrum at species level in children in Yili

To accurately characterize bifidobacteria in school-age (pre-school) children in Yili, Xinjiang, we selected the *groEL* gene for high-throughput sequencing to amplify intestinal bifidobacteria and developed a comprehensive analysis pipeline for bifidobacteria species/subspecies (see methods). Following this pipeline, we obtained a total of 9,257 representative ZOTU sequences, of which 6,958 were identified as bifidobacteria following species annotation. While non-bifidobacteria ZOTUs constitute a substantial proportion (24.83%) of the count, they represented a mere 2.34% of the abundance. The amplification results of the 16S rRNA gene analysis revealed that high-throughput sequencing based on the *groEL* gene accounted for only 5.69% of unclassified *Bifidobacterium*, which is significantly lower than the 28.86% observed for the 16S rRNA gene. The *groEL* gene identified 22 species/subspecies of bifidobacteria, whereas the 16S rRNA gene identified only 11 species. It is noteworthy that the *groEL* gene was capable of distinguishing bifidobacteria at the subspecies level, while the 16S rRNA gene could only differentiate them at the species level. With regard to relative abundance, significant differences were observed for *B. adolescentis*, *B. bifidum*, *B. catenulatum*, and *B. pseudocatenulatum* (Table S4).

The analysis revealed that *B. adolescentis* (36.32%) and *B. longum* subsp. *longum* (19.24%) were the most abundant bifidobacteria in children from Yining, Xinjiang. Additionally, *B. pseudocatenulatum* (8.95%) and *B. breve* (9.41%) were also identified. This study further identified several bifidobacteria with very low abundances (relative abundance less than 0.01%), including *B. animalis* subsp. *animalis*, *B. longum* subsp. *infantis*, *B. longum* subsp. *suillum*, *B. crudilactis*, *B. merycicum*, *B. longum* subsp. *mongoliense*, and *B. gallinarum*. Notably, none of these extremely low-abundance bifidobacteria were detected by the 16S rRNA gene.

Furthermore, to demonstrate the accuracy of Bifidobacteria classification, we integrated our bifidobacterial sequences with 30 *groEL* genes from NCBI that represent model bifidobacterium sequences to construct phylogenetic trees (Fig. 5a). The analysis of the bifidobacterial phylogenetic tree demonstrated that the representative ZOTU sequences derived from the intestinal tract of children in Yining, Xinjiang, at the same species or subspecies level clustered within the same evolutionary clade as their corresponding reference type strains, closely resembling these reference type strains. In addition, the analysis revealed that different species of bifidobacterium occupy distinct evolutionary branches in their phylogeny and are phylogenetically distant from one another.

**Figure 5.**
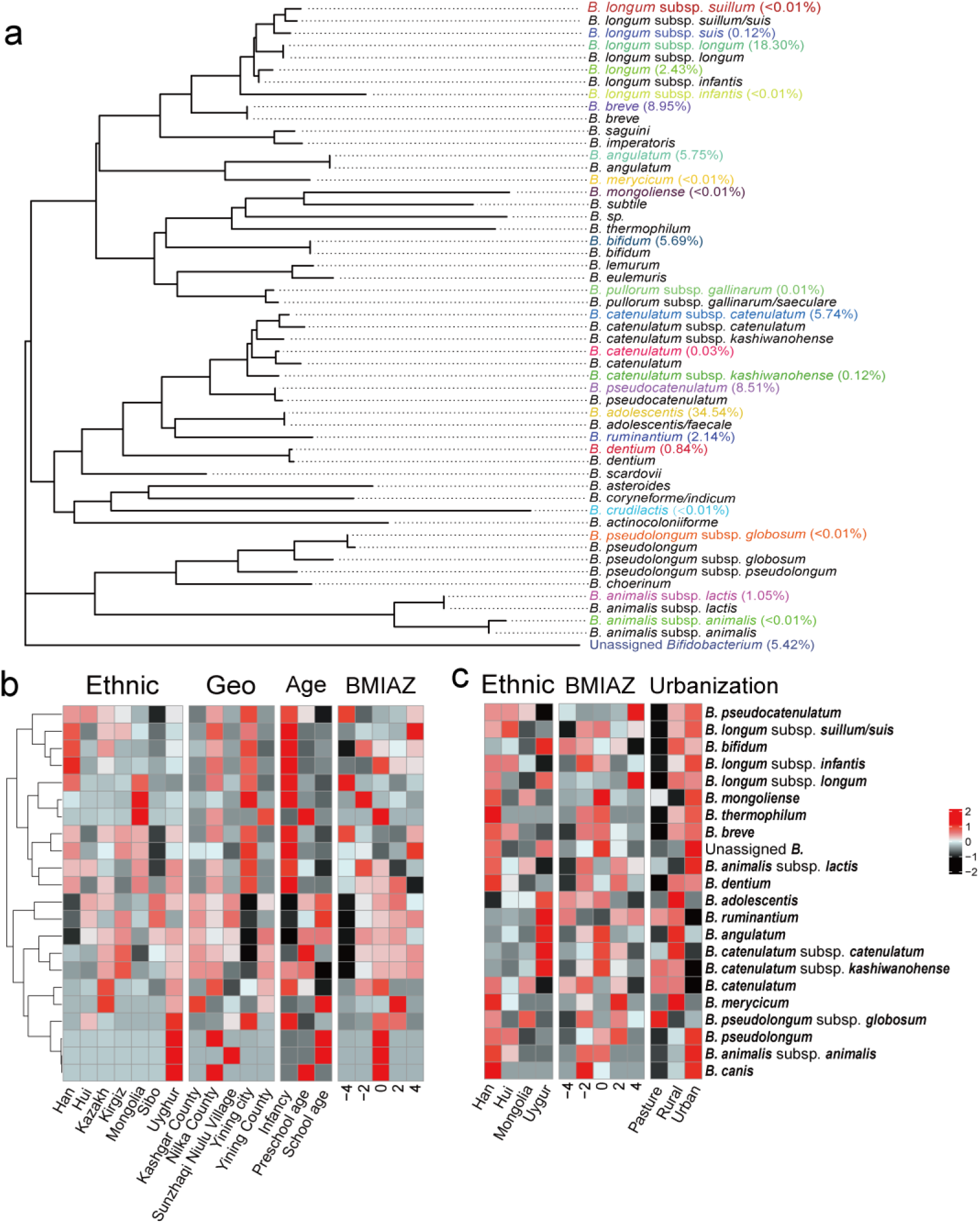
The abundance and composition of *Bifidobacterium*. a) Phylogenetic relationship at species level of *Bifidobacterium.* Phylogenetic tree constructed using the *groEL* gene nucleotide sequences, employing the generalized time-reversible (GTR) model for analysis. The phylogenetic tree was constructed by selecting representative amplicon sequence variants (ZOTUs) (denoted in color) at each species or subspecies level, determined by the most abundant ZOTUs for each taxon, and combined with 30 *groEL* gene sequences (depicted in black) sourced from the complete genome assemblies of representative *Bifidobacterium* available in the NCBI database. b) This study investigates the associations between ethnicity, geography, age, and BMI factors with the abundance of *Bifidobacterium* species. c) Lu study investigates the associations between ethnicity, BMI and urbanization factors with the abundance of *Bifidobacterium* species. Heat map showing the mean abundance determined by normalization at each of the factors. (This study, n = 776; Lu, n = 394.)

### Gut Bifidobacterium distribution and result reproducibility

To characterize the associations between the Bifidobacterial community and important factors, including geography, age, ethnicity and BMI, we analyzed the correlations between the normalized relative abundance of each covariate. In addition, to assess the reproducibility of the Bifidobacterium distribution, we incorporated an independent and comparable dataset, referred to as the “Lu study.” From this dataset, we extracted a subset of samples comprising children aged 0 to 18 years, with a significant proportion of Han Chinese participants. The heatmap depicted the distribution of 22 abundant species, highlighting a significant enrichment of *Bifidobacterium longum* subsp. *infantis* in the Han population, while *Bifidobacterium mongoliense* was notably enriched in the Mongolian population (Fig. 5b). This finding was corroborated by similar results obtained in the “Lu study” (Fig. 5c). It is noteworthy that our infant samples were collected from two locations - the Yining City and the Nilka County. Importantly, the urban area (Yining City) only had infant samples. After accounting for the potential confounding effects of different grouping variables, it was found that *B. catenulatum* and *B. catenulatum* subsp. *Kashiwanohense* were absent from the urban infant samples. This observation is consistent with previous studies, such as the “Lu study”, which have identified these Bifidobacterium species as being associated with pasture or rural lifestyle people.

### Bifidobacterium changes in the developmental trajectory of children

To investigate the dynamic changes in Bifidobacterium across the developmental trajectory of children, we calculated the mean relative abundance of bifidobacterial communities of adjacent ages. The relationship between the logarithmic relative abundance and age of 14 *Bifidobacterium* species/subspecies with a relative abundance exceeding 1% was analyzed using a Loess regression model (Fig. 6). The analysis revealed that *B. adolescentis*, *B. angulatum*, and *B. ruminantium* exhibited an initial increase in relative abundance with age, followed by stabilization. In contrast, *B. dentium*, *B. longum*, and *B. breve* demonstrated a decline in relative abundance with age before reaching a stable phase, while *B. longum subsp. longum* maintained a consistently stable relationship with age throughout. Additionally, *B. bifidum* and *B. catenulatum subsp. catenulatum* displayed fluctuating patterns in their association with age. Furthermore, the observed trends in bifidobacterial dynamics were largely consistent with those reported in the “Lu study” (Fig. S5). However, it is noteworthy that the stabilization of bifidobacterial development in our study occurred at approximately 6 years of age, whereas in the “Lu study”, this stabilization was observed at around 4 years of age, indicating a relatively slower developmental trajectory in our findings.

**Figure 6.**
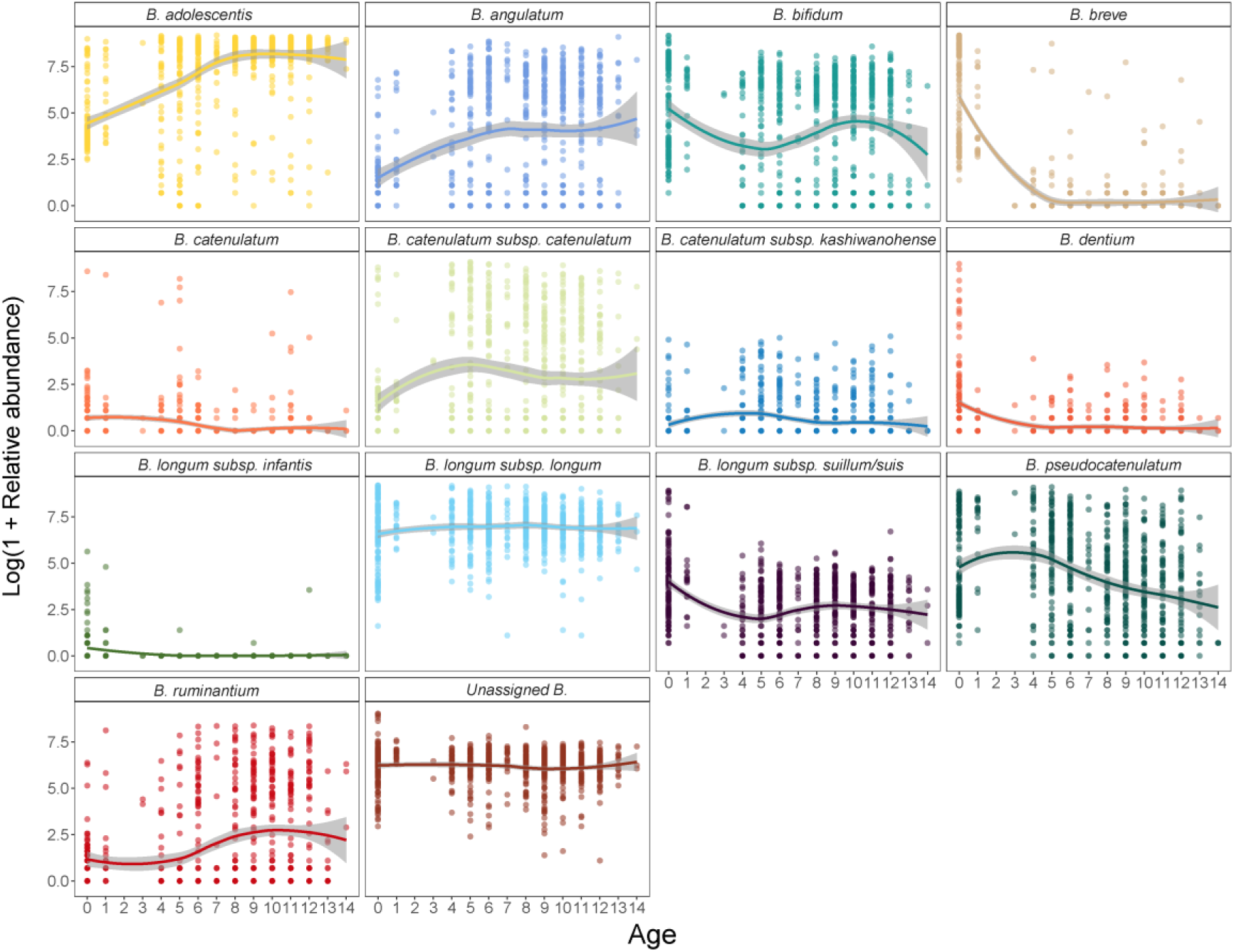
Association between Age and differential *Bifidobacterium* species. Linear regression analysis was performed for different *Bifidobacterium* species/subspecies using a loose model, and different colors represented different *Bifidobacterium* species/subspecies.

### Consistency of ZOTU effects across different studies and enthnic groups

Given the limitation of the present sample set to regional children and the inclusion of multiple ethnic minority groups from China, it is essential to investigate the consistency across different ethnic groups. First, we identified suggestively significant ZOTUs using the negative binomial distribution (ZINB) model and subsequently evaluated the effect sizes of these signals across diverse datasets and their respective ethnic subgroups. The significant effect sizes related to developmental trajectories in our dataset showed poor correlation with the estimated effects from the “Lu dataset” (Fig. 7 and Fig. S6). Subsequent to this, we applied Deming regression (see Methods) to more effectively account for measurement error and evaluate calibration. We then merged the “Lu dataset” into a unified dataset referred to as “ALL” to calculate the overall effect size. Although effect sizes were highly correlated across studies, notable divergences were observed. Specifically, the regression slope was 1.53 for Ili versus ALL (95% CI: 0.72–1.08), 0.74 for the Lu versus ALL (95% CI: 0.71-0.77) and 0.25 for the Lu versus ALL (95% CI: 0.24-0.26) (Fig. 7a). In addition, we compared the deviations in effect sizes across different ethnic groups within their respective datasets. For instance, the Uyghur population showed no deviation in the Ili samples (β_Deming_ = 0.99, 95% CI: 0.96-1.01), whereas the Kazakh population (β_Deming_ = 1.61, 95% CI: 1.14-2.09) exhibited a shift. A similar pattern was observed that the Han population demonstrated an offset in the “Lu study” (β_Deming_ = 1.25, 95% CI: 0.87-1.63) (Fig. 7b). Subsequently, we then compared the shifts in effect sizes among the three representative ethnic groups: Uyghur, Kazakh, and Han. Notably, both Uyghur and Kazakh exhibited substantial deviations from the Han (β_Deming_ = 4.40, 95% CI: 4.12-4.68; β_Deming_ = 4.33, 95% CI: 3.76-4.89), while a high degree of consistency was observed between Uyghur and Kazakh (β_Deming_ = 1.08, 95% CI: 0.99-1.17) (Fig. 7c).

**Figure 7.**
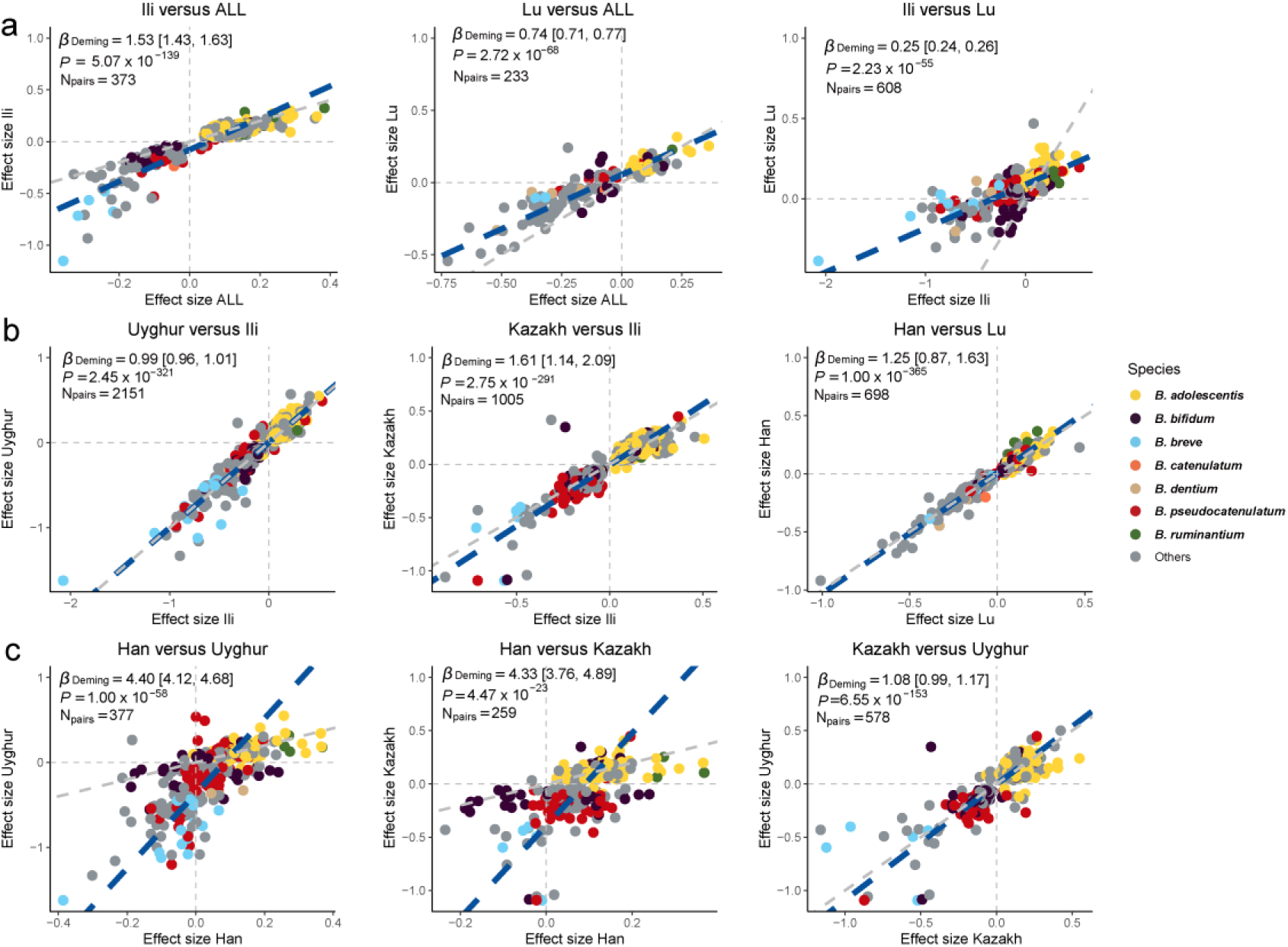
Effect sizes of ZOTUs for age correlate between different studies and ethnic groups. The ZINB model was employed to estimate the effect size of each ZOTU on age. Scatterplots with the effect sizes from our study and the “Lu Study”. a) Estimates of ZOTUs effect sizes were compared between our study and Lu study. b) Estimates the effect sizes of ZOTUs in each study compared with the ethnic groups within the study. c) Estimates of ZOTUs effect sizes were compared among different ethnic groups.

**Figure 8.**
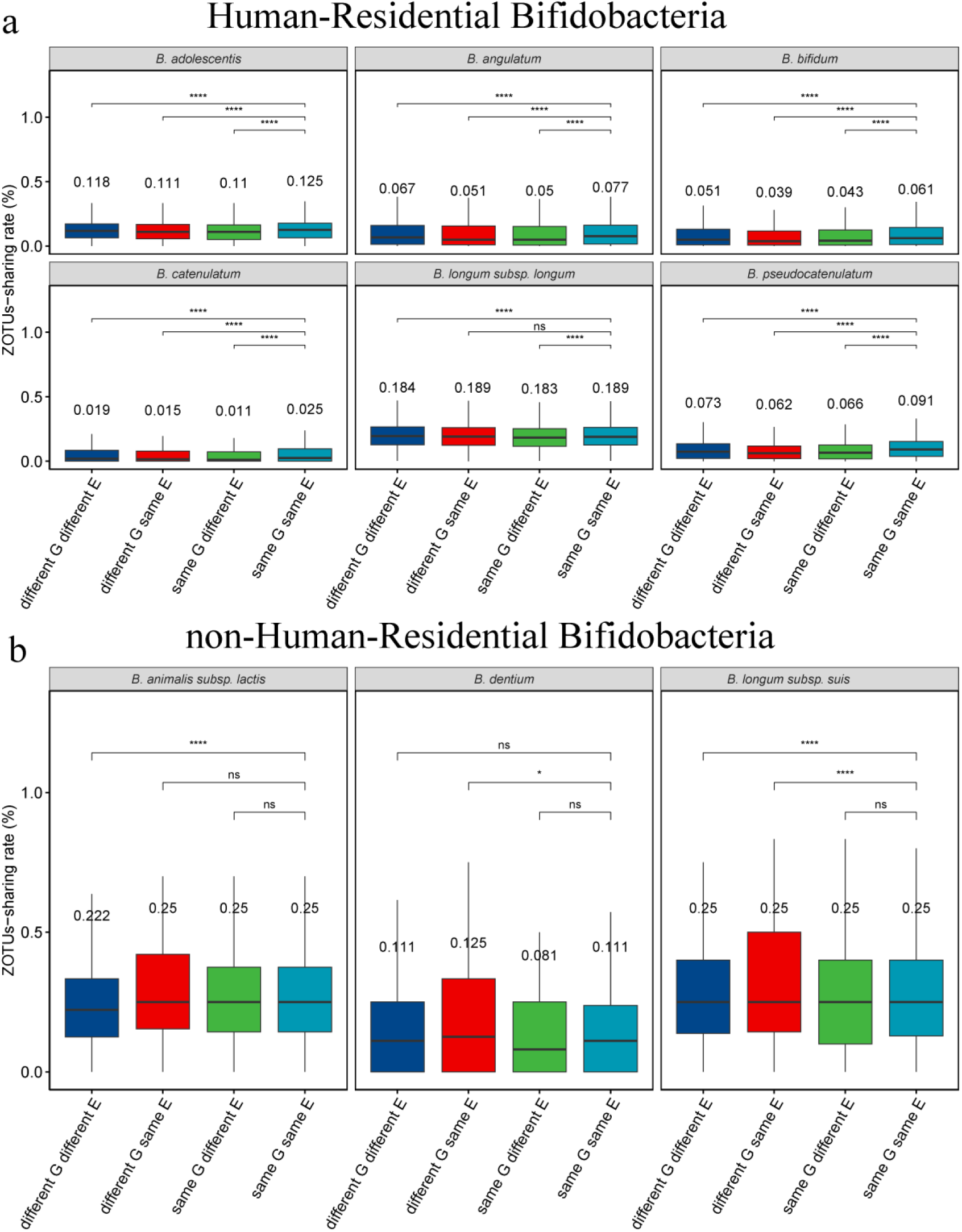
Children ZOTUs-sharing rates (number of shared ZOTUs / total number of unique ZOTUs × 100%) across different ethnic and regional types. In the box plots, distinct colors represent different categories: G for geographic and E for ethnic. The term “same G same E” refers to the sharing rate of two individual ZOTUs among children from the same geographic location and belonging to the same ethnic group. Conversely, “different G same E” indicates the ZOTUs sharing rate in pairwise comparisons among children of the same ethnic group residing in different geographical locations. All comparisons yielded as determined by the Kruskal-Wallis test. Select bifidobacterial species of human origin as: a) human-residential bifidobacteria (HRB); and b) non-HRB, which includes species that are natural inhabitants of animals or the environment.

Broadly, our findings provide evidence that the effect sizes for Bifidobacterium ZOTUs exhibit reasonable consistency between the Ili and other studies, justifying further trans-ethnic approaches to improve discovery power in developmental trajectory of children association studies. Moreover, these analyses underscore the variability in the influence of intestinal bifidobacteria on aging across different ethnic populations, highlighting the inconsistency of this effect among diverse groups. Given the current sample size, the analysis of the growth and developmental trajectories of children influenced by intestinal bifidobacteria across different ethnic groups remains inconclusive, underscoring the necessity for further investigation in future studies.

### Transethnic and Transgeographical Gut Bifidobacterium Sharing Rates

The colonization of bifidobacteria is established during the newborn period (mother-to-child transmission(41)), suggesting that its transmission is unlikely to occur via social interactions among children. Furthermore, bifidobacteria may serve as biomarkers for growth and development in children, with potential variations influenced by ethnic backgrounds. To support this perspective, and in accordance with the guidance provided by Wong,(42) bifidobacteria were classified into two categories: human-residential bifidobacteria (six Species) and non-human residential bifidobacteria (three Species). A comparative analysis was conducted of transethnic and transgeographical sharing rates of these species and subspecies of bifidobacteria among children. The analysis focused on four classifications: 1) the same area with different ethnic groups, 2) the same ethnic groups across different regions, 3) the same area with varying ethnic groups, and 4) different ethnic groups in diverse regions. The conclusions drawn from this analysis are as follows: Nearly all human-residential bifidobacteria exhibited significantly higher Sharing Rates within the same ethnic group residing in the same location, in comparison to other classifications (*P* < 0.001); But in non-human bifidobacteria, *B. dentium*, *B. animalis subsp. lactis*, and *B. longum subsp. suis* exhibited no significant correlation with either ethnic or regional effects; consequently, no discernible pattern was observed.

## Discussion

The gut microbiota is crucial for human health, particularly as bifidobacteria are considered important commensal microbes and probiotics within the children gut microbiome. Differences in microbiota between ethnicities may serve as a mediator of health disparities.(43, 44) However, there is a paucity of studies investigating gut microbiota among different ethnicities in specific regions, particularly during childhood. In this study, we recruited 776 infants and school-aged children from Yining, Xinjiang. We performed amplicon sequencing targeting the 16S rRNA gene and the *groEL* gene to systematically analyze the diversity, composition, influencing factors, and potential functions of intestinal microorganisms in children. Additionally, we identified characteristic marker bacteria associated with the growth processes of children in this region.

Body mass index is a crucial indicator of a child’s growth and development. It is subject to rapid fluctuations and does not exhibit a straightforward linear relationship with age.(45) However, different methods and measures can lead to different inferences about children’s health.(46) Furthermore, due to unbalanced regional development(47) and ethnic specificities(48), the early growth and development trajectories may vary, and child health surveys focusing on minority populations in remote regions are generally inadequate.(49, 50) In accordance with the standards established by the World Health Organization (WHO), we developed a standard normal deviation statistical index (Z-score) for the intestinal microbiota of children from specific ethnic groups. This index was utilized to quantitatively differentiate and assess the growth standards of healthy children. There are significant differences in the growth and development of children from various ethnic groups (Fig. S1). However, there are limitations, as we lack an ethnic-specific body index reference dataset.

We divided the sample into three age groups: infants, preschool-aged children, and school-aged children. The results revealed that *Bifidobacterium* serves as the dominant intestinal core microbe in these children which differs markedly from that observed in the global population.(37) The alpha diversity and microbial composition in infants was lower than that observed in both school-aged and preschool-aged groups. At the same age, children from different sampling sites did not exhibit significant differences in gut microbiota composition. The analysis of various species across different developmental periods revealed that *Bifidobacterium*, *Streptococcus*, and *Enterococcus* were significantly associated with the infant group. These genera are classified as amphioanaerobic or anaerobic and play a crucial role in early life colonization.(51) Furthermore, significant differences were observed in the composition of intestinal flora among children of varying ages, particularly in the abundance distribution of *Actinobacteria*, *Firmicutes*, *Proteobacteria*, and *Bacteroidetes*. Functional predictive analyses indicated that the gut microbiota of healthy preadolescent children exhibited greater gene enrichment (approximately 6% of gene families) and distinct functional pathway distributions.(52) This finding suggests that the developmental trajectory of the gut microbiome progresses toward increased maturity and functional diversity as children are exposed to new environments.(53) The reduction of *Bifidobacterium* and the introduction of new microorganisms may have further facilitated this transition, providing valuable insights into how human microbiomes evolve in conjunction with host matures.(54, 55)

To describe the characteristics and biomarkers of maturation of intestinal microflora in (pre) school-age children. We selected pediatric samples with BMI Z-scores ranging from -2 to 2 to calculate “relative microbiota maturity”.(39) The results showed that there was no significant difference in microbiota maturity of children with different health conditions. This is inconsistent with previous findings.(56-58) This instead indicates that BMI alone is insufficient for assessing the malnutrition or obesity of children, not only because of the rapid growth but also the influence of other diseases related to abnormal gut microbiome development.(59) Consider the complexity of microbiome count data (zero-inflation and over-dispersion, etc), we construct a ZINB model to effectively accommodate these characteristics. The results identified a cluster of ZOTU taxa exhibiting a significant negative correlation with age, while phylogenetic analyses indicated that these taxa were primarily affiliated with *B. breve*. At the same time, we also observed a slightly significant positive association between *B. adolescentis* and age, which may also be due to their ability to degrade plant-derived carbohydrates and tend to become more dominant in adulthood.(60)

These findings underscore the critical role of bifidobacteria in the growth and development of children. To conduct a comprehensive analysis of bifidobacteria, we developed a bifidobacteria-specific amplification process based on high-throughput sequencing (see the follow-up report for details) to accurately annotate bifidobacterial species at the species level. The results are exciting, demonstrating superior performance of our method in both the detection of low-abundance (less than 0.01%) and accurate annotation compared to 16S rRNA sequencing. We observe that higher relative abundances of specific Bifidobacterium species (e.g., *B. catenulatum* and *B. catenulatum* subsp. *kashiwanohense*) are associated with individuals’ pasture or rural lifestyle.(16, 61)

Additionally, we incorporated another study, referred to as the “Lu study” In terms of host developmental trajectories, our findings indicate that stabilization of bifidobacterial development occurred at approximately 6 years of age, whereas the “Lu study” reported this stabilization at around 4 years of age, suggesting a relatively slower developmental trajectory in our results. Furthermore, we compared our study with the “Lu study” to examine the heterogeneity of bifidobacteria in relation to age benefits across different ethnic groups. We do observe heterogeneity across ethnic groups (Uyghur vs Kazakh, β_Deming_ = 4.40, 95% CI: 4.12-4.68). This is consistent with previous hypothesis that intestinal microbial development in children of different ethnic groups is inconsistent.(62) Finally, we conducted a study on transethnic and transgeographical sharing rates of gut bifidobacteria. human-residential bifidobacteria (HRB) show a high degree of specificity with ethnic groups, while non-HRB exhibit no significant correlation with either ethnic or regional effects. Certain non-HRB, such as *B. animalis* subsp. *lactis*, are significantly prevalent in dairy products(63-65) compared to other bifidobacteria species, but do not dominate the human gut.

Our study reveals the association between microbiome dynamics and child growth trajectories, highlighting the importance of bifidobacteria. However, it is crucial to acknowledge not only the complex developmental variables but also the heterogeneity associated with ethnicity. Recent literature indicates that ethnicity is linked to various factors, including diet, lifestyle, socioeconomic status, and systemic issues like racism.(66, 67) Although we have captured some rapid changes in child growth, it is possible that not all causative factors have been identified. This may be attributed to limitations in sequencing techniques and the size of the sample. Last but not least, in the precise screening, design, and application of probiotic bifidobacteria, it is essential to consider the uneven development observed among different ethnic groups.

## Materials and Methods

### Study design

We performed an observational study of fecal microbes in preschool and school-age children across ethnic groups and regions, with the approval of the Science and Technology Ethics Committee for the First Affiliated Hospital of Shihezi University (approval number 2017-117-01) according to the ethical guidelines outlined in the Declaration of Helsinki and its amendments.(68, 69) This study aims to investigate the gut microbial markers associated with the developmental trajectories of children and how these trends may interact with ethnic and geographic factors. We have received active cooperation and strong support from various schools and families.

During the early stages of study design, we communicated with the authorities, community service centers, primary school principals and kindergarten principals of five regions (Kashgar County, Nilka County, Sunzhaqi Niulu Village, Yining County and Yining City) of Ili Kazak Autonomous Prefecture in Xinjiang, China, to get permission. Subsequently, we provided training for teachers and staff from the Women’s Federation, who then communicated the purpose and significance of our research to students and their families. Informed consent forms were distributed to students to take home, allowing parents to decide whether to sign them. Additionally, parents of preschoolers were also given the opportunity to consent to their children’s participation in the study.

Our work entails a protracted and systematic process. Despite the presence of numerous ethnic groups in Xinjiang, the samples we collected predominantly consisted of Uyghur and Kazakh populations, while the samples from other ethnic groups were relatively few. We will continue to expand the sample size in the future.

### Sample collection

In our research, we first investigated which schools or regions belong to multi-ethnic communities:

For school-aged children: we communicated with local authorities and school principals to get permission to sample. In addition, the researchers trained the teachers, and the teachers who then publicized the purpose and significance of our research. At the same time, informed consent forms are issued, which the students take home for parents to decide whether to sign or not. The following days, the participants had their height and weight measured and were inquired about their date of birth and recent dietary habits. After the survey, the participants were supplied with a stool sampler, an ice bag, and an aseptic bag, and were given along with comprehensive guidance on how to collect and preserve the samples. Participants are required to send their stool samples back to the school as soon as possible, ideally within 1 to 2 days.

For infants: we collaborate with the Fourth Division Hospital of the Xinjiang Production and Construction Corps. After the hospital staff informs the expectant mothers about the purpose of our study, those who agree to participate provide their contact information. Subsequently, we conduct door-to-door sampling visits after the infants are born. The method of sample collection employed is consistent with the procedures previously mentioned.

Inclusion criteria: 1). school-aged children; 2). Ethnic minorities, and both parents are ethnic minorities; 3). Able to provide informed consent; 4). No antibiotics or other medications that could affect the composition of gut microbes have been used in the past 6 months; 5). Be willing to provide regular stool samples as required by the study. Exclusion criteria: 1). Individuals suffering from severe digestive disorders (e.g. Crohn’s disease, ulcerative colitis) or other chronic diseases (e.g., diabetes, heart disease); 2). Recent (within 6 months) use of antibiotics or other drugs that affect the gut microbiota; 3). Individuals who are unable to provide informed consent.

All the collected samples were temporarily stored in the portable vehicle refrigerator at -20°C and transported back to the laboratory (Shihezi University Food Biotechnology Research Center) within 48 h and was stored at -80°C until DNA extraction. The sampling and transportation process touches the environment as little as possible, and the sampler wears sterile gloves throughout this process.

### DNA extraction and sequencing

As in previous study,(40) the samples were thoroughly homogenized with sterile phosphate-buffered saline (PBS) and the stool homogenization procedures were conducted within a biosafety cabinet, completed within 30 min to ensure sample integrity and minimize contamination risks. Approximately 1 mL of the resulting mixture was subsequently aliquoted for DNA extraction. The total genomic DNA was then extracted from the fecal samples using the E.Z.N.A. Soil DNA Kit (Omega Biotek, Norcross, GA, U.S.) according to manufacturer’s instructions. After the quantitative test of DNA purity met the standard using NanoDrop2000, the DNA was transported on dry ice and the V1-V3 region of the bacterial 16S rRNA gene and the *groEL* gene of the *Bifidobacterium* functional gene were sequenced on the Illumina Novaseq 6000 (Illumina Inc., San Diego, CA, USA) of Majorbio Bio-Pharm Technology Co., Ltd (Shanghai, China).

### 16s rRNA sequencing and analysis pipeline

Primer 27F-YM (5’-AGR GTT YGA TYM TGG CTC AG-3’) and 534R (5’-ATT ACC GCG GCT GCT GG-3’)(70) were used for amplicon sequencing in the V1-V3 region of the bacterial 16S rRNA gene and the PCR conditions were as follows: start at 95℃ for 4 min, 30 cycles denaturing at 95℃ for 30 s, annealing at 72℃ for 50 s, and final extension at 72℃ for 10 min. Referring to the EasyAmplicon pipeline developed by Liu Yongxin,(71) the raw paired-end sequence data (Raw reads data) was processed using “ebarcode” (https://github.com/jianhuang525/ebarcode) to remove the adapters and primers from the raw data. Meanwhile, a sliding window strategy was adopted to trim the low-quality bases at the ends. Appropriate trimming of the low-quality bases at the ends can significantly improve the success rate of paired-end sequence merging. The “vsearch”(72) software was employed with the “--fastq_mergepairs” command to merge the paired-end reads. Quality control was carried out using the “-fastx_filter” command (with parameters selected as -fastq_maxee_rate 0.005 and -fastq_minlen 400) to obtain high-quality reads. Subsequently, the “--fastx_uniques” command was further utilized to merge the unique reads, resulting in a non-redundant sequence set. After that, the sequences were first arranged according to sequence abundance using “--uchime_deno” and self-dechimerization was performed. Then, the “--uchime_ref” command was applied, combined with the silva_138.1_ssu(73) database, to conduct dechimerization again. The sequences after dechimerization were clustered into Amplicon Sequence Variants (ZOTU)(74) using the “--cluster_unoise” command. For the clustered ZOTU sequences, the “sintax”(75) algorithm was used to align with the rdp_16s_v19_sp(76) database for species annotation (with parameters selected as: -strand = both, -sintax_cutoff = 0.6). The “otutab_filter_nonBac.R” script was used to remove chloroplasts, mitochondria, etc. The “--usearch_global” command was employed to align the quality-controlled sequences to generate the original feature table (with identity = 0.97). Finally, the R package “vegan” was utilized to rarefy the sequence counts of each sample to a standardized total of 10,000 sequences.

### groEL gene sequencing and analysis pipeline

The extracted DNA was also used for sequencing the functional *groEL* gene of *Bifidobacterium*. Bifidobacteria-specific primers Bif-Groel-F (5’-TCC GAT TAC GAY CGY GAG AAG CT-3’) and Bif-Groel-R (5’-CSG CYT CGG TSG TCA GGA ACA G-3’) were used for amplification and the corresponding conditions for the amplification strategies were predenaturation at 95℃ for 5 min; 35 cycles of denaturation at 95℃ for 45 s, annealing at 60℃ for 45 s and extension at 72℃ for 60 s were repeated. The annotated database utilized and the upstream analysis workflow can be accessed at https://github.com/jianhuang525/groELpipeline. This process is currently in the testing phase, as detailed in a forthcoming article.

### Evaluation criteria of body mass index of children

We calculated the z-scores for weight, height, and body mass index (BMI; kg/m²) standardized by age and sex using the zscorer package (v.0.3.1)(77) in the R software environment (v4.3.0). This software strictly adheres to the World Health Organization (WHO) standards for child growth. According to the standards, 567 individuals (81.3%) fell within the normal range (-2 to 2sd), while 106 individuals (15.2%) were classified as short (< -2 sd), 24 individuals (3.4%) in the high range (> 2 sd). According to the WHO-reported trends in height changes by age, girls tend to be taller than boys until approximately 13 years of age. However, after 13 years, boys surpass girls in height, and this difference remains stable into adulthood. During the early stages of infancy (0-2 years), growth is relatively rapid. Between 3 and 13 years of age, growth stabilizes, and height exhibits a linear relationship with age.

### Regression model

We employed both the random forest algorithm and the zero-inflated negative binomial (ZINB) model to evaluate effect sizes for each taxon.

Random Forests regression model:

Random Forest regression analysis was performed using the default settings of the R implementation (R package ‘randomForest’). Specifically, the model utilized 10,000 trees (ntree = 10,000) and employed the default value for mtry, calculated as *p*/3, where *p* represents the number of input ZOTUs(78). Intestinal microbiome samples were obtained from healthy individuals, typically those with a BMI Z-score ranging between -2 and 2. Operational taxonomic units (ZOTUs) with fewer than 10 counts per sample were excluded from the analysis. Modeling regression was conducted to examine the relationship between the relative abundance of ZOTUs and the actual age of each child at the time of fecal sample collection, employing a Random Forest machine learning algorithm. The significance of the model fit was evaluated by comparing it to a null model, in which the age labels of the samples were randomly permuted relative to their 16S rRNA gene microbiome profiles (*P* < 0.0001, based on 9,999 permutations). A ranked list of bacterial taxa was generated by identifying those taxa whose relative abundance contributed the most to the mean square error when excluded. The predictive performance of the model was further assessed as a function of the number of top-ranked taxa, based on their feature importance scores.

Zero-inflated Negative Binomial (ZINB) regression model:

The complexity of microbiome count data, especially in scenarios characterized by inflation and overdispersion, hinders us from elucidating the associations between microorganisms and host phenotypes. Bayesian zero-inflated negative binomial (ZINB) regression model that is capable of identifying differentially abundant taxa associated with distinctive host phenotypes and quantifying the effects of covariates on these taxa(40). Prior to analyses, ZOTUs with exceptionally low abundance (fewer than 30 observed counts across the cohort) were excluded, adhering to the suggestion of Wadsworth.(79) To facilitate convergence in the analysis of covariate information, the age variable was centered, and the BMI Z-score was utilized. Employing a Bayesian zero-inflated negative binomial (ZINB) regression model, we systematically analyzed each of the 9,219 taxa (ZOTUs) individually, accounting for the effects of ethnicity and geography, as well as their interaction.

### The “Lu study”

In a similar study on *groEL* gene sequencing involving 1,674 participants, we selected 394 children’s samples and processed the original sequencing data using the same analytical approach as this study. After quality control, 314 samples remained for further analysis. To compare phenome-wide effect sizes across different groups, we computed Pearson correlation estimates, thereby quantifying the correlation between age-relative taxon effect sizes from the “Lu study” samples and our samples, as well as among different ethnicities. To account for any errors in the effect estimates, we employed Deming regression(80), a form of error-in-variables total least squares regression effect sizes, utilizing the deming() function from the R package deming (version 1.5).

### Statistical analysis

All statistical analyses were conducted using the R software environment (version 4.3.0). The analyses of microbial alpha and beta diversity, along with differential analysis, were performed using the “vegan”(81) and “microeco”(82) packages within the R. To compare the abundances of microbial taxa and functional profiles, we employed non-parametric tests: the two-sided Wilcoxon rank-sum test for pairwise comparisons and the Kruskal-Wallis test for comparisons involving more than two groups, unless otherwise specified. We analyzed the predicted functional profiles of microbial communities using the PICRUSt2(83) software (version 2.5.0). The data representing functional KO orthologs from the KEGG ORTHOLOGY database for each sample were utilized for further analysis. These results were analyzed using the ggpicrust2(84) package, employing the ‘pathway_daa’ function with the DESeq2 method and applying Benjamini-Hochberg corrections for multiple comparisons.

## Data Availability

The raw amplicon sequence data reported in this study have been deposited in the Genome Sequence Archive in National Genomics Data Center, China National Center for Bioinformation / Beijing Institute of Genomics, Chinese Academy of Sciences, 16S rRNA (GSA: CRA022029) and *groEL* gene (GSA: CRA022030) that are publicly accessible at https://ngdc.cncb.ac.cn/gsa.

## Author contributions

Yongqing Ni, Jian Huang and Yuxin Chu conceived and designed the study. The supervision and guidance of the project was provided by Yongqing Ni and Fengwei Tian. Yuxin Chu, Quanhao Zhao and Huimin Zhang contributed to the collection and organisation of samples and data. Jian Huang performed the statistical analyses and interpreted the findings. Jian Huang, Yuxin Chu and Zhixuan Liang wrote the manuscript. All authors contributed to the article and approved the submitted manuscript.

## Acknowledgments

The authors would like to thank the anonymous reviewers for the help in improving the manuscript.

## Statement of Ethics

The studies involving human participants were reviewed and approved by the Ethics Committee of the First Affiliated Hospital, Shihezi University School of Medicine (2017-117-01). The participants provided their written informed consent to participate in this study.

